# The maize hexokinase gene *ZmHXK7* confers salt resistance in transgenic *Arabidopsis* plants

**DOI:** 10.1101/2021.04.26.441367

**Authors:** Qianqian Liu, Zengyuan Tian, Yuqi Guo

## Abstract

The hexokinase (HXK) gene family, whose members play vital roles in sugar induction signals and glycolysis in organisms, is widely found in plants. Although some hexokinase genes have been studied in maize, a systematic report of the gene family and its role in plant resistance is lacking. In this study, 10 hexokinase genes were systematically identified in maize based on the maize genome-wide database. Phylogenetic analysis divides the maize HXK protein family into four clusters. Prediction of cis-regulatory elements showed that a number of elements responding to abiotic stress exist in the promoter of hexokinase genes. The expression profile of these genes, originated from B73, showed that different members of hexokinase genes are highly expressed in roots and leaves of maize under salt or drought stress, which is similar to that of Mo17.The coding sequence of *ZmHXK7* gene, isolated from maize B73, was constructed into plant expression vector pMDC45 and then transformed into *athxk3* (Salk_022188C). By hyg resistance detection, PCR analysis, and western blot confirmation, the homozygous progenies of transgenic Arabidopsis lines were identified. Subcellular localization analysis showed that the *ZmHXK7* gene was located in cytosol. Seedling growth and salt stress inhibition in complementary mutant plants of *ZmHXK7* gene were significantly improved, and enhanced salt tolerance was displayed. Our study provides insights into the evolution and expression patterns of the hexokinase gene and show that maize ZmHXK7 proteins play an important role in resisting salt stress, which will be useful in plant breeding for abiotic stress resistance.

## 1 Introduction

The hexokinase (HXK) protein, which promotes hexose phosphorylation and plays a large role in sensing and conducting sugar signals as a bifunctional enzyme, was first discovered in yeast bread (Meyerhof 1927), and the wheat (*Triticum aestivum* L.) hexokinase gene was first isolated and identified from higher plants (Saltman 1953). It is a starter enzyme for glucose glycolysis in living organisms, and is also the rate-limiting enzyme in the glycolytic pathway (Olsson et al. 2003; Perata et al. 1997). The high expression of tomato *LeHXK2* in flowers is not accompanied by the increase of HXK enzyme activity, indicating that hexokinase not only catalyzes respiration metabolism (Menu et al. 2001), but also has the function of sugar perception (Karve et al. 2010). The regulatory role of HXK2, a yeast glucose phosphorylating enzyme, as a glucose sensor is independent of its metabolic role. Evidence for this is provided by a *hxk2* mutant protein with reduced catalytic activity but which is still fully functional in glucose signaling (Mayordomo and Sanz 2001).

Previous studies showed that the plant HXK gene exists in multi-gene families (Karve et al. 2010); for example, rice has 10 family members (Cho et al. 2006), Arabidopsis thaliana has 6 (Karve et al. 2008)and tomato (*Lycopersicon, escule*) has 4 (Kandel-Kfir et al. 2006). Most HXK members of the gene family in some plants are forecast to contain 9 exons and mainly distribute in mitochondria, and a few members exist in cytoplasm, chloroplast and plasmalemma (Rajagopal Balasubramanian and Physiology 2007). Different subcellular localization of plant HXK, which is closely related to its gene structure, lead to functional differences (Karve et al. 2008). Based on phylogenetic analysis,HXK proteins can be broadly divided into two categories (Cho et al. 2006; Giese et al. 2005; Kandel-Kfir et al. 2006; Olsson et al. 2003),one with plastid signal peptide structure and the other with N-terminal membrane anchoring (N-terminal membrane anchors) structure (Damari-Weissler et al. 2006; Kandel-Kfir et al. 2006; Karve et al. 2010; Kim et al. 2006) which determine the mitochondrial localization of HXK. A fraction of the AtHXK1 protein associated with mitochondria can translocate to the nucleus where it acts as a glucose sensor to control gene transcription (Cho et al. 2010). HXK activity located on the mitochondria is strongly inhibited in the presence of ADP, n-phthalein glucosamine and mannose ketone (Bruggeman et al. 2015; Cheng et al. 2011; Graham et al. 2007), which is different from activity on other organelles. HXK located in chloroplast can also be divided into two categories. One kind of HXK is located in the chlorophyl, and its function is to phosphorylate hexose. Another type of HXK binds to the chloroplast membrane to phosphorylate glucose transferred from the chloroplast. HXK distributed in cytoplasmic matrix may only exist in monocotyledon plants such as rice and maize (Nilsson et al. 2011). and small bowl thistle (Karve et al. 2010). *OsHXK4* is a chloroplast matrix HXK, which is the first HXK of this type found in monocotyledons (da-Silva et al. 2001), while *OsHXK7* is a cytoplasmic HXK (Cho et al. 2006).

In human tumor tissue, hexokinase, especially hexokinase II (HXKII), activity is increased (Shulga et al. 2009). Plant HXK is reported to respond to environmental changes and interact with nutrients, light and excitation signals to form a network that controls the growth and development of *Arabidopsis thaliana* (Dai et al. 1999; Kim et al. 2013; Moore et al. 2003). HXK and ethylene interact closely and mediate signal transduction pathways (Zhou et al. 1998). Glycosylase-mediated sugar signaling requires a complete ABA signal transduction chain (Huijser 2000). Both hypoxia and cold stress can stimulate the increase of hexokinase activity in plant roots (Fox et al. 1998). The increased activity of HXK in guard cells will accelerate the closure of stomata, thereby reducing transpiration and reducing water loss in plants. Stomatal closure is induced by sugar and mediated by abscisic acid (Yu et al. 2013). In addition to the above functions, *Arabidopsis AtHKLl* has been found to be a negative regulator of plant growth (Karve and Moore 2009) and can regulate anthocyanin synthesis (Hosokawa et al. 1996; Kawabata et al. 1995; Minakuchi et al. 2008; Solfanelli 2006; Vitrac et al. 2000; Zheng et al. 2009). Therefore, HXK is closely related to abiotic environment regulation by interacting with hormones in plants.

In a previous study, nine members of the ZmHXK gene family were identified (Galina et al. 1999). Multiple sequence alignment analysis showed that maize ZmHXK protein may be involved in plant hormone and abiotic stress response, glucose inhibition, light and circadian rhythm regulation, Ca^2+^ response, seed germination and transcriptional activation of CO_2_ response (Zhang et al. 2014). While *ZmHXK5* and *ZmHXK6* were characterized, little is known about detailed functions of *ZmHXK7* gene which may be involved in abiotic stress response. In this study, a genome-wide identification of hexokinase gene was performed in inbred lines B73 and Mo17. The phylogenetic relationship, conserved domains, genome organization, and gene structure of hexokinase were systematically studied. The CDS sequence of *ZmHXK7* was cloned and constructed into the plant expression vector pMDC45, and then mutant *athxk3*(Salk_022188C) was infected with Agrobacterium tumefaciens. T3 transgenic plants *athxk3* 355S:*ZmHXK7* (AT1G47840) were obtained. Seed germination, plant growth and salt tolerance were analyzed, which provide essential information concerning the physiological functions of ZmHXK and important theoretical basis for genetic engineering in maize salt tolerance enhancement under salt stress.

## 2 Materials and methods

### 2.1 Sequence analysis and identification of maize HXK gene family

cDNA sequence, coding sequence, protein sequence, and promoter sequences (upstream 2,000 base of transcriptional starting site) of maize HXK family genes were downloaded from Genomics Network (http://plants.ensembl.org/index.html). Mapchart2.2 (http://mg2c.iask.in/mg2c_v2.0/) was used to map the relative loci of the maize HXK genes according to the length of each chromosome and the position of each member on the chromosomes.

### 2.2 Phylogenetic, gene structure, and conserved domain analysis of hexokinase genes

MEGA7.0 multisequence comparative analysis of amino acids was performed to construct the phylogenetic tree of hexokinase proteins in *Zea mays, Arabidopsis,Oryza sativa*, and *Sorghum* using the neighbor-joining algorithm with 1000 bootstrap replications. Gene structure was obtained by the online program Gene Structure Display Server (http://gsds.cbi.pku.edu.cn/) using hexokinase cDNA and genome sequences. The online website Multiple Em for Motif Elicitation (MEME, http://meme-suite.org/tools/meme) was used to identify conserved domains of hexokinase proteins.

### 2.3 Analysis of promoter, prediction of subcellular localization, prediction of three-dimensional modeling, and interacting networks of hexokinase proteins in Oryza sativa and Arabidopsis

Around 2000bp of genomic DNA sequence upstream of the promoter of hexokinase genes were selected for analysis of cis element through online website plant CARE (http://bioinformatics.psb.ugent.be/webtools/plantcare/html/). Online website ProtComp9.0 (http://linux1.softberry.com/berry.phtml?Topic=protcomppl&group=proloc) was used to predict subcellular localization of ZmHXK.

The online network ExPASy (https://web.expasy.org/protparam/) was used to predict molecular weight and isoelectric point. Ident and Sim (http://www.bioinformatics.org/sms2/ident_sim.html) was used to carry out sequence homology comparison analysis. The online website SingnalP4.1 (http://www.cbs.dtu.dk/services/SignalP-4.1/) was used to forecast signal peptide. PfamScan (https://www.ebi.ac.uk/Tools/pfa/pfamscan/) was used to query the function of the conservative base sequence. GraphPad. Prismv5.0 was used to map the relative expression of genes.

Three-dimensional modeling of Hexokinase proteins was performed by Phyre2 server http://www.sbg.bio.ic.ac.uk/~phyre2/html/page.cgi?id=index). The prediction of the interacting networks of proteins was constructed in STRING (https://string-db.org/?Tdsourcetag=s_pctim_aiom).

### 2.4 Plant Material and abiotic stress treatment

The cultivated maize (B73 and Mo17) seedlings were used to detect the expression of hexokinase gene family. Seeds were selected and sterilized on the surface, then were sown in pots filled with mixed substrate of vermiculite and perlite in a 3:1 ratio, and were placed in plastic trays (3 pots per tray). Seeds germinated and were cultivated for experiments. The three-leaf stage were selected for drought and salt stress treatment, with three replications for each treatment. Control plants were irrigated with 1/2 Hoagland nutrient solution daily. Salt stress was applied by watering the plants in the same way but using solutions supplemented with NaCl to final concentrations of 0.5% or 1%. The top second leaves and roots at 0,2,6,24,72 h after salt stress were harvested for RNA extraction. Moderate and severe drought stress was performed by withholding irrigation of the pots for 2 weeks: after this period, the control was watered with half of the nutrient solution once every two days. 1/2 Hoagland nutrient solution was irrigated in moderate drought treatment, and 1/2 Hoagland nutrient solution was irrigated in drought treatment After severe drought stress, the seedlings were rewatered. The top second leaves and roots of the samples were used for RNA extraction.

### 2.5 RNA Extraction and Quantitative RT-PCR

The total RNA in treated root and leaf of maize was extracted by Invitrogen’s TRizol Reagent. The extracted RNA was reverse-transcribed into cDNA using the Kangwei century HIFI-MMLV cDNA first strand synthesis kit, and the primers were designed in online tools and synthesized by ShangYa biotechnology co., LTD. Primer sequence is shown in table 1. Real-time fluorescence quantitative reaction was carried out in Roche lighCycler480 using Kangwei century fluorescence quantitative PCR kit (Ultra SYBR Mixture (LOW ROX)); the reaction system was 20uL, including 10uL SYBR Green Master MIX (2X), 10uM gene-specific primers, 1.0uLcDNA and 7.0uL ddH20. The 2 ^−ΔΔct^ method was used for data calculation, and the relative gene expression was represented by mean ± standard deviation. The heat map was drawn for tissue expression profiling by online website (http://www.omicshare.com).

**Table 1.**
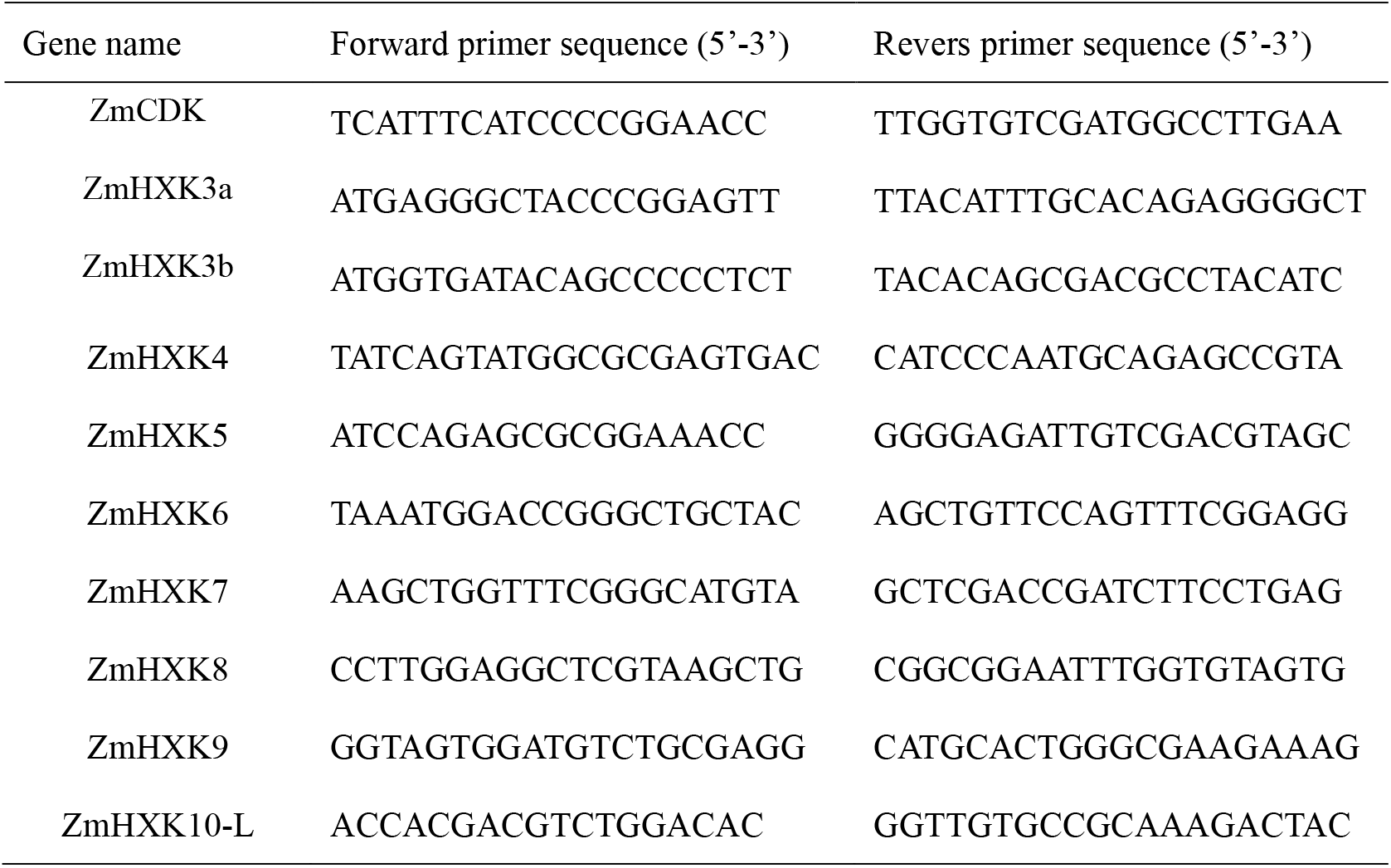
Primer sequences used in quantitative real-time PCR analysis of ZMHXKs

### 2.6 Construction of plant expression vector and transformation of A.thaliana

The cDNA of *ZmHXK7* with open reading frame was obtained by PCR using maize total RNA as template. The *ZmHXK7* gene was amplified by PCR using the forward (5’-CACCATGGAGAAGCAAGGCGTCGAC −3’) and reverse primers (5’-GTACTGCGAGTGGGCAGCTGCAATG −3’), which are specific for *ZmHXK7* gene. PCR was performed using standard conditions consisting of initial denaturation at 94°C for 4 min followed by 35 cycles of 94°C for 30s, 56°C for 30s and 72°C for 2 min, followed by a final extension of 72°C for 5 min. The PCR products of *ZmHXK7* were purified using Kangwei century Gel Extraction kit (CW23025) and used to construction entry vector (Invitrogen, USA) by TOPO cloning reaction according to the manufacturer’s instructions. An entry vector containing the gene of the correct orientation and sequence was used to construct the target vector pMDC45, using LR ClonaseII enzyme mediated gateway cloning reaction according to manufacturer’s protocol. The recombinant plasmid pMDC45 was introduced into *A*.*tumefaciens* strains GV3101 by liquid nitrogen freeze-thaw method. The *A*.*tumefaciens* transformation was performed using floral dipping technique. Transformed Arabidopsis plants were grown in greenhouse under a 16/8 h light/dark cycle at 24/22 °C with 70% relative humidity. Seeds harvested from the transformed plants (T0) were grown on MS medium containing 25 mg L^−1^ hyg under the same growth conditions. Homozygous T3 progeny including *athxk3* 35S:*ZmHXK7* derived from T2 population were selected and confirmed by PCR for further salt tolerance analysis.

In accordance with the T-DNA Primer Design (http://signal.salk.edu/tdnaprimers.2.html), we set up two paired reactions, LP+RP and LB+RP (*athxk3*-FP: 5’-TTGTTGAAACGGTTGATCTTCC-3; *athxk3*-RP: 5’-TGGCAATTCTGGAGAAG ATTG-3’; LB: 5’-ATTTTGCCGATTTCGGAAC-3’) to identify homozygous mutants.

### 2.7 Confirmation of transgenic Arabidopsis plant

Genomic DNA of T3 transgenic Arabidopsis was extracted from the leaves of transgenic and wild type seedings that grew for 6 weeks using the CTAB method. PCR was performed using primers GFP −FP (5’-ATGAGTAAAGGAGAAGAACTTTTCAC TGGA-3’) and *ZmHXK7* (5’-GTACTGCGAGTGGGCAGCTGCAATG −3’), and the amplification was performed using the program: 94 °C for 2 min, followed by 35 cycles of program (94 °C for 30 s, Temperature °C for 30 s, 72 °C for 2 min), and terminated by an extension at 72 °C for 5 min. The PCR products were detected by electrophoresis on 1.5% agarose gel.

### 2.8 Western blot analysis of transgenic Arabidopsis plant

WT and transgenic Arabidopsis plants were grown on MS medium containing 25mg L^−1^ hyg. After 10 days of cultivation, total proteins of the seedlings were extracted from WT and transgenic Arabidopsis seedings with a buffer consisting of 50 mM Tris/HCl (pH 8.0), 150 mM NaCl, 1 mM EDTA, and 0.2% (w/v) Triton X-100, 4% β-mercaptoethanol, 1 mM dithiothreitol (DTT) and 1% (v/v) protease inhibitor cocktail and then used for protein quantification with BCA protein quantitative Kit (Abcam ab6789). The protein sample (200μg amounts) were electrophoresed in 8% SDS-PAGE and the gels were transferred to nitrocellulose membranes. The membranes were blocked with TBST buffer (10 mM Tris/HCl, pH 7.5, 150mM NaCl and 0.05% Tween-20) supplemented with 5% non-fat milk for 2 h and incubated with primary antibodies (Anti-GFP antibody, abcam290, GAPDH Monoclonal Antibody abcam38689 diluted at 1:1000) in TBST buffer with 5% BSA overnight at 4 °C. Afterwards, the membranes were washed three times (10 min each) with TBST buffer and incubated with the secondary antibodies (Goat Anti-Rabbit IgG H&L(HRP)Abcam6721,Goat Anti-Mouse IgG H&L (HRP), abcam6789, dilution at 1:1000) for 2 h. After three washes with TBST buffer, the membranes were incubated with a <u>chromogenic</u> agent Enhanced HRP-DAB Chromogenic Substrate Kit (Boster).

### 2.9 Subcellular localization assay

Fluorescence of ZmHXK7-GFP fusion proteins in transgenic plants was observed using confocal laser-scanning microscopes. The roots from 4-weeks-old transgenic Arabidopsis plants were selected to be observed.

### 2.10 Determination of hexokinase activity (HK)

The hexokinase (HK) activity assay kit (BC0740) purchased from Solarbio was used to detect the content of hexokinase with WT, *athxk3* and *athxk3* 35S::*ZmHXK7*. The production of 1nmolNADPH per minute per g of tissue is defined as a unit of enzyme activity.

### 2.11 Salt stress treatments involving the transgenic Arabidopsis plant

The seeds of WT, *athxk3*, and *athxk3* 35S::ZmHXK7 were disinfected with sodium hypochlorite. The seeds were planted on MS medium under aseptic conditions, vernalized at 4°C for 3 days, and cultured in a light incubator for 6 days. The seeds of WT, *athxk3*, and *athxk3* 35S::*ZmHXK7* of the same size were moved to the following MS plates in Ultra-clean table: 0mM NaCl, 0mM glucose;150mM NaCl, 0mM glucose;0mM NaCl, 100mM glucose;150mM NaCl, 100mM glucose. Physiological indices such as dry and fresh weight were measured after 6 days in the light incubator.

### 2.12 Determination of chlorophyll content

Chlorophyll was extracted with ethanol as solvent. 0.1g leaves were soaked in 25 mL 90% ethanol and extracted overnight (16 h). The sample supernatant was measured at 663 nm and 645 nm using UV-1800PC spectrophotometer (Arnon 1949).

The calculation formula of chlorophyll content is as follows (mg/g):

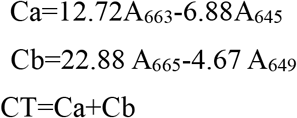

A663: light absorption value at 663 nm; A645: light absorption value at 645nm; Ca: concentration of chlorophyll a; Cb: concentration of chlorophyll b; CT: Total concentration of chlorophyll and chloroplast pigment content = (CT X extraction liquid volume)/ fresh weight of the sample

### 2.13 Determination of proline content

Leaf and root samples were homogenized with 5mL 3% thiosalicylic acid, centrifuged for 10min at 12000r/min. 2 mL glacial acetic acid and 2 mL acidic ninhydrin were added into 2.0 mL supernatant. After mixing, the mixture was heated in a water bath at 100°C for 60 minutes. After cooling, the mixture was extracted with 4ml toluene. Taking toluene as blank, the absorbance was measured at 520nm. According to the regression equation below, the content of proline (X μg/2 mL) in 2mL determination solution was calculated, and then the percentage of proline content in the sample was calculated (Pinheiro and Bates 1995).

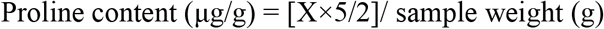

### 2.14 Determination of malondialdehyde content

Weave 0.1g of plant leaves, add a small amount of quartz sand into the mortar, add 2ml 10% trichloroacetic acid, grind until homogenate, then add 3ml 10% trichloroacetic acid for further grinding, and centrifuge homogenate at 4000r/min for 10min. Add 2ml supernatant to the test tube, add 2ml distilled water to the tube, and then add 2ml 0.6% thiobarbituric acid solution to each tube. Shake well, react in boiling water bath for 15min, then cool quickly and centrifuge. The absorbance values of supernatants were measured at 532 nm and 450nm respectively (Zhang et al. 2008).

Malondialdehyde content calculation:

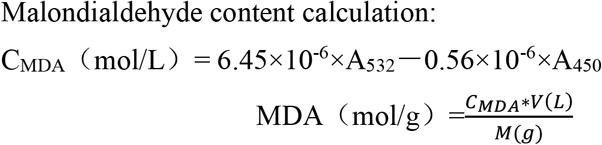

### 2.15 Statistical Analysis

GraphPad Prism 5 was utilized for all statistical analysis Samples were analyzed in triplicates, and the data are expressed as the mean ± standard deviation unless noted otherwise. P<0.05 was considered significant (*), P<0.01 was considered very significant (**) and P<0.001 was considered extremely significant (***).

## 3 Results

### 3.1 Identification of Members of Maize Hexokinase Gene Family

To identify the Hexokinase gene family in maize, online website Ensembl plant (http://plants.ensembl.org/index.html) was used to search members of HXK gene families. A total of 10 hexokinase genes in maize were identified (table 2), its genetic sequence information displayed in table 2. The length of coding region of HXK family genes in maize ranged from 609-1530bp. The molecular weight and isoelectric point were calculated using ExPASy (https://web.expasy.org/protparam/). Our results showed that the deduced protein sequences of hexokinase gene family members ranged from 202 to 509 amino acids, among which the protein encoded by *ZmHXK4* is the longest (509aa); The molecular weight of ZmHXK protein ranged from 23 to 55KDa, and the molecular weight of *ZmHXK10L* is the smallest(23KDa). The isoelectric points of all ZmHXK protein ranged from 5.13 (*ZmHXK8*) to 9.12 (*ZmHXK10L*), among which only *ZmHXK10L* (9.12) were alkaline, the rest were acidic. Ten HXK genes of maize were distributed on 4 of the 10 chromosomes according to database, among which there were four genes (*ZmHXK3a, ZmHXK6, ZmHXK8* and *ZmHXK9*) located on chromosome 3, two genes (*ZmHXK3b, ZmHXK4*) on chromosome 8, three genes (*ZmHXK5, ZmHXK7, ZmHXK10*) on chromosome 6. And only one gene (*ZmHXK10L*) on chromosome 9.

**Table 2.** Sequence information of maize HXK gene family

| Gene | Gene ID | CDs(bp) | Chromosomal location(bp) | Protein length(aa) | Molecular weight(kDa) | pI | Subcellular localization | Signal peptides |
| --- | --- | --- | --- | --- | --- | --- | --- | --- |
| ZmHXK3a | Zm00001d042146 | 1494 | Chr.3:153564761-153570791 | 497 | 53.50 | 5.60 | Cytosol/<br>Mitochondrial | - |
| ZmHXK3b | Zm00001d011889 | 1494 | Chr.8:164038137-164038137 | 497 | 53.41 | 5.81 | Cytosol/<br>Mitochondrial | - |
| ZmHXK4 | Zm00001d010796 | 1530 | Chr.8:128039610-128045077 | 509 | 54.86 | 6.18 | Cytosol/<br>Mitochondrial | - |
| ZmHXK5 | Zm00001d038792 | 1524 | Chr.6:164389095-164395441 | 507 | 54.69 | 6.15 | Cytosol/<br>Mitochondrial | - |
| ZmHXK6 | Zm00001d043511 | 1521 | Chr.3:202355009-202361480 | 506 | 55.04 | 5.92 | Cytosol/<br>Mitochondrial | - |
| ZmHXK7 | Zm00001d037689 | 1380 | Chr.6:134063370-134066401 | 459 | 49.3 | 5.14 | Cytosol/<br>Mitochondrial | - |
| ZmHXK8 | Zm00001d039512 | 1512 | Chr.3:6248365-6252317 | 503 | 54.37 | 5.13 | Cytosol/<br>Mitochondrial | - |
| ZmHXK9 | Zm00001d043607 | 1515 | Chr3:204995530-205005583 | 504 | 54.98 | 5.93 | Cytosol/<br>Mitochondrial | - |
| ZmHXK10 | Zm00001d037861 | 1512 | Chr.6:140078685-140083862 | 503 | 54.24 | 5.58 | Cytosol/<br>Mitochondrial | - |
| ZmHXK10L | Zm00001d046061 | 669 | Chr.9:60696813-60697811 | 222 | 23.89 | 9.12 | Mitochondrial | - |

Based on results of subcellular localization prediction with CELLO (http://cello.life.nctu.edu.tw/) and WOLF PSORT (https://wolfpsort.hgc.jp/), *ZmHXK10L* is predicted to locate only to the mitochondria, the rest ZmHXK in the cytoplasm or mitochondria. Signal peptide prediction using SingalP - 4.1 (http://www.cbs.dtu.dk/services/SignalP-4.1/) showed that corn HXK gene families had no signal peptide.

### 3.2 Phylogenetic and Evolutionary Analysis of Hexokinase Gene Family

A phylogenetic tree was constructed to assess the evolutionary relationship of hexokinase proteins among *Zea mays, Arabidopsis, Oryza sativa, and Sorghum (* Fig 1) by using Clustal Omage (https://www.ebi.ac.uk/Tools/msa/clustalo/).The thirty-five hexokinase proteins from four different species were classified into four distinct clusters (clusters I to IV). The ZmHXK gene was present in all of the four clusters.

**Fig. 1.**
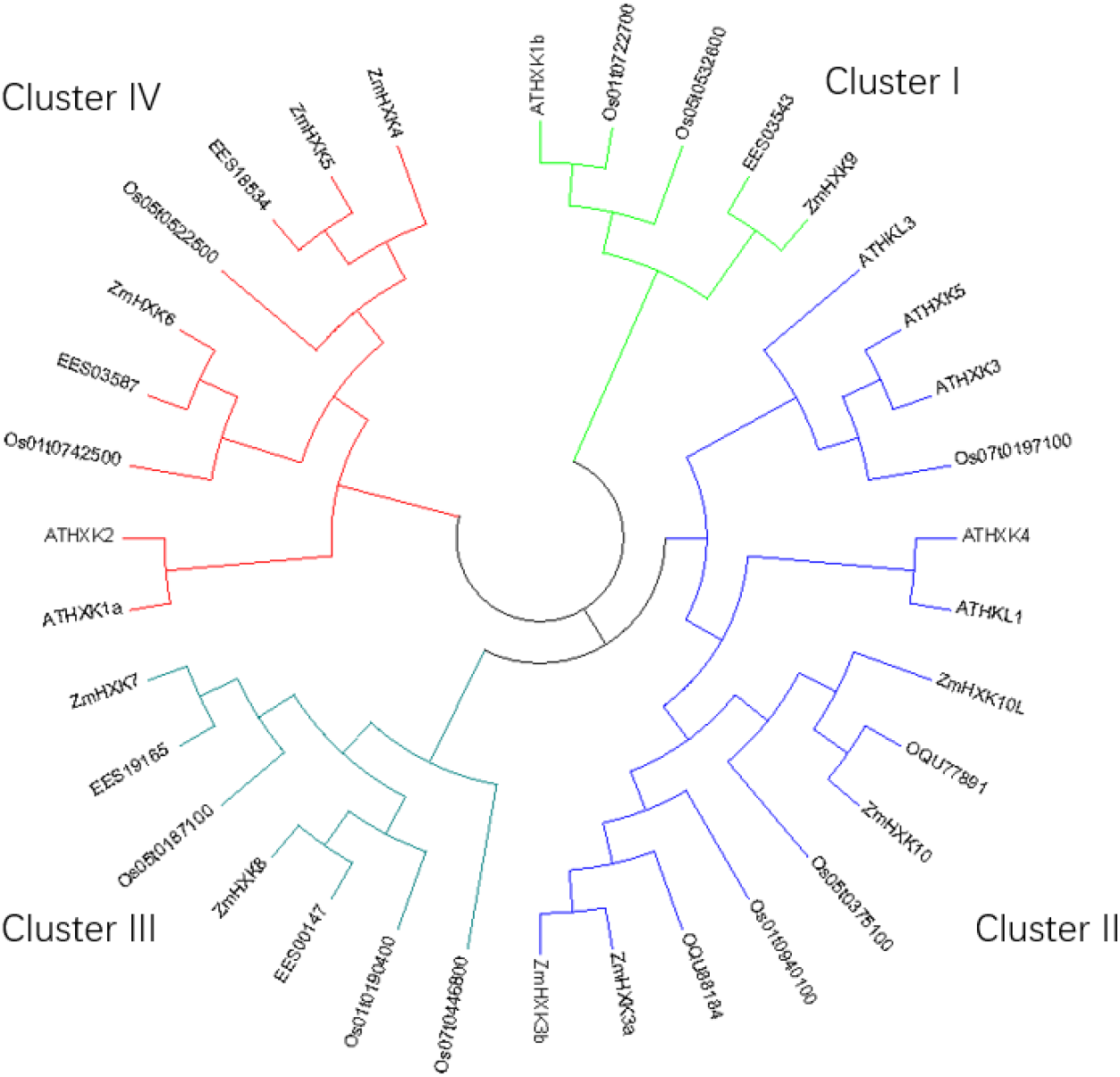
Phylogenetic analysis of hexokinase proteins from *Zea mays, Arabidopsis, Oryza sativa, and Sorghum*.

*ZmHXK4, ZmHXK5* and *ZmHXK6* are paralogous genes in cluster IV; *ZmHXK7* and *ZmHXK8* were grouped into cluster III, which include all hexokinase genes originated from monocotyledons. These results show that these genes were evolutionarily conserved within monocotyledons, which provides evidence for the functional research of the uncharacterized but evolutionarily conserved genes.

### 3.3 Genomic Structure, Conserved Domain, and Motif Analysis of Hexokinase Proteins in Maize

The genetic structure of ZmHXK family members showed that the hexokinase genes contain 9 exons, except for *ZmHXK7* and *ZmHXK10L*, which contains 7 and 3 exons respectively (Fig.2). Predicted motif (table 3) analysis revealed that 7 of 10 members of hexokinase proteins contained all 10 motifs (Fig.2). Two genes (*ZmHXK7* and *ZmHXK8*) had no motif 1. One gene (*ZmHXK10L*) only had 4 motifs, resulting in incomplete *ZmHXK10L* protein. Distribution features of the 10 predicted motifs in deduced proteins were consistent with the phylogenetic analysis.

**Fig. 2.**
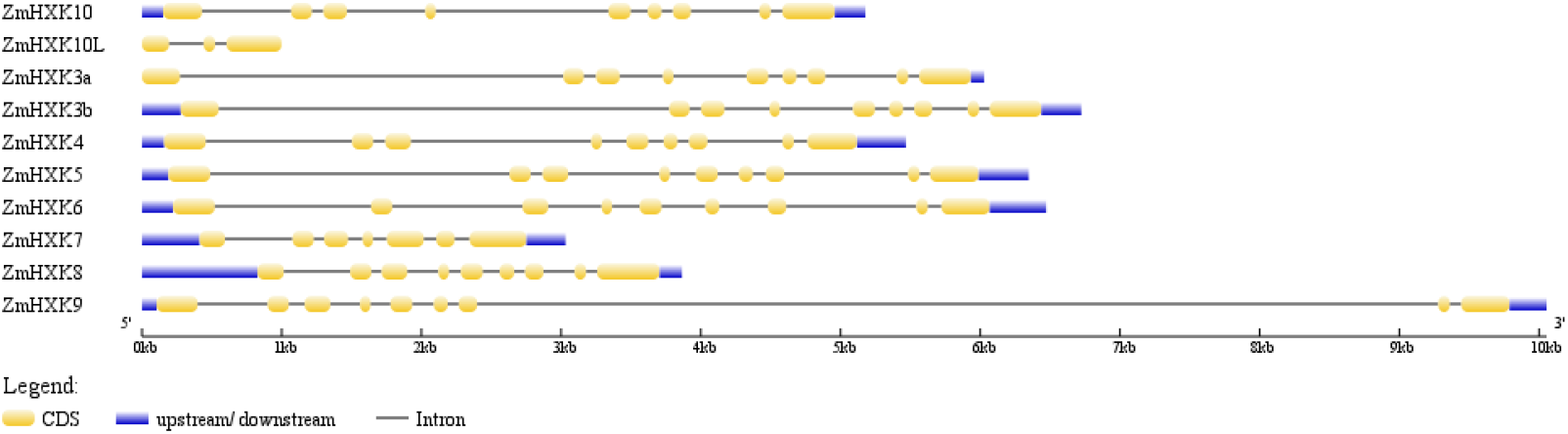
HXK gene structure of maize

**Fig. 3.**
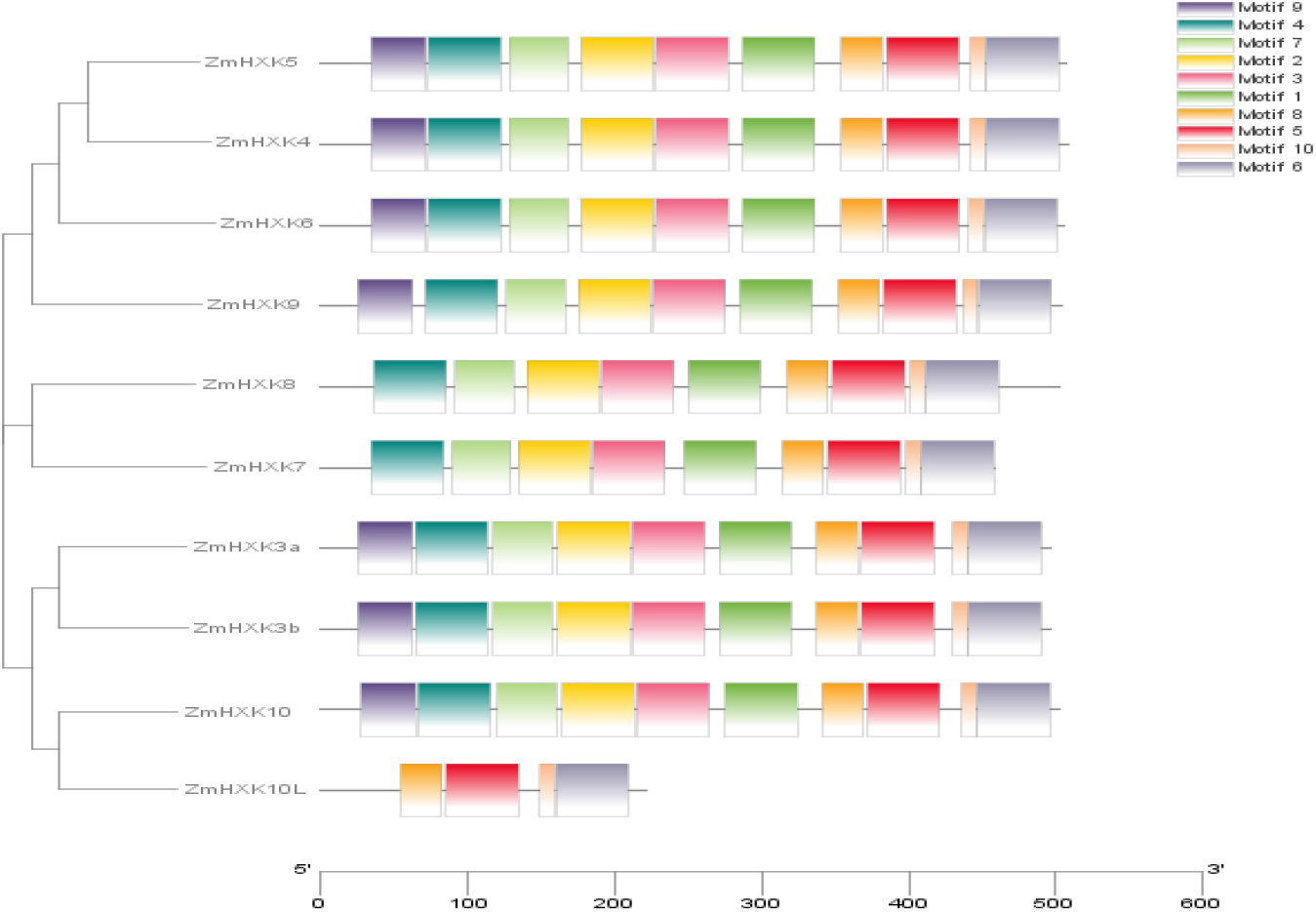
HXK conservative sequence analysis of maize

**Table 3.**
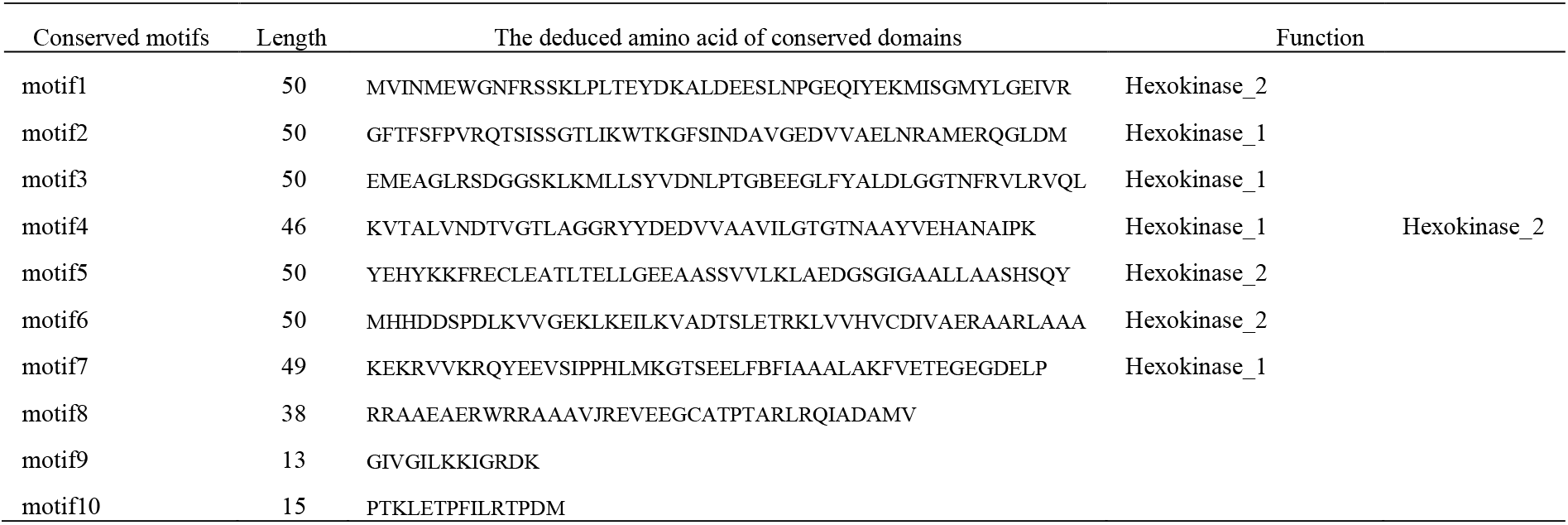
HXK conservative motif analysis of maize

The PfamScan database was used to query functions of motif; results show that motif 1-motif 7 has known functional records in the database, among which motif 2, 3, 4 and 7 encode *AtHXK1* domain, while 1,5 and 6 encode *AtHXK2*. Notably, motif 4 was predicted to encode both HXK1 and HXK2 domain of Arabidopsis (Table 3).

### 3.4 Promoter analysis of the hexokinase gene family in maize

To assess the possible response patterns of the hexokinase gene to different abiotic stress treatments, cis-acting elements were predicted in the promoter region of hexokinase gene family members using Plant CARE database.

Our results show that the promoter region of all HXK gene members contains different plant hormone response elements (Fig.4). ABRE (abscisic acid response element), G-box, MYB and STRE is distributed in the promoter region of all ten ZmHXK members of family. Methyl jasmonic acid TGACG-motif is distributed in nine ZmHXK genes except for *ZmHXK10*. AuxRR (auxin response element), EREE (thylene response elements), GARE (gibberellin) and salicylic acid TCA response element is distributed in promoter region of 4, 3, 4, and 4 HXK members of ZmHXK family, respectively. In addition, each HXK gene promoter region also contains different numbers of components responding to abiotic stress response including anaerobic stress response components ARE and GC-motif (enhancer-like element involved in anoxic specific inducibility), DRE (Dehydration-Responsive-Element), LTR (Low Temperature Response) and stress-inducible or defense-related response element TC-rich repeats. Ten HXK genes contain different numbers of multiple cis-acting elements related to plant growth or light response, such as G-box, box 4, GA-motif, GT1-motif, MRE, SP1. Some cis-acting elements such as CAT-box related to meristem, GCN4 involved in endosperm expression and O2-site related in zein metabolism regulation appear in the promoter of some ZmHXK genes of hexokinase family. These data suggest that ZmHXK may be involved in the response to abiotic stresses such as drought and salt stress, anaerobic, hypothermia as well as to hormones regulation.

**Fig. 4.**
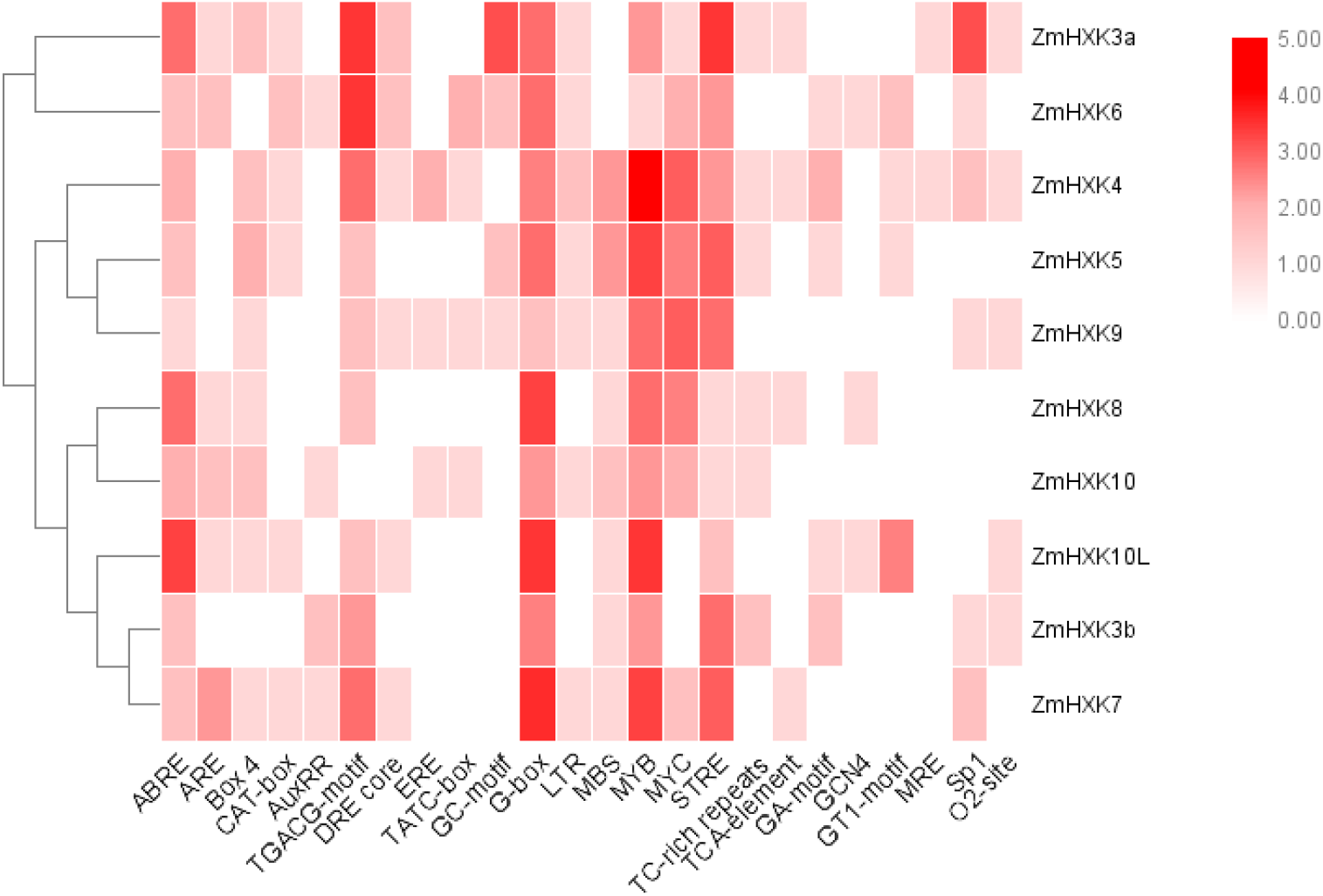
analysis of cis-acting elements in promoter region of maize HXK gene family

### 3.5 Prediction of Three-Dimensional Modeling and Interaction of Hexokinase Proteins in maize and Oryza sativa

To understand the structural characteristics and interaction network of the Hexokinase subfamily, three-dimensional models of the all the Hexokinase proteins were constructed (Fig.5), and the protein-protein interactions network in Oryza sativa and maize were predicted using STRING(Fig.6).

**Fig. 5.**
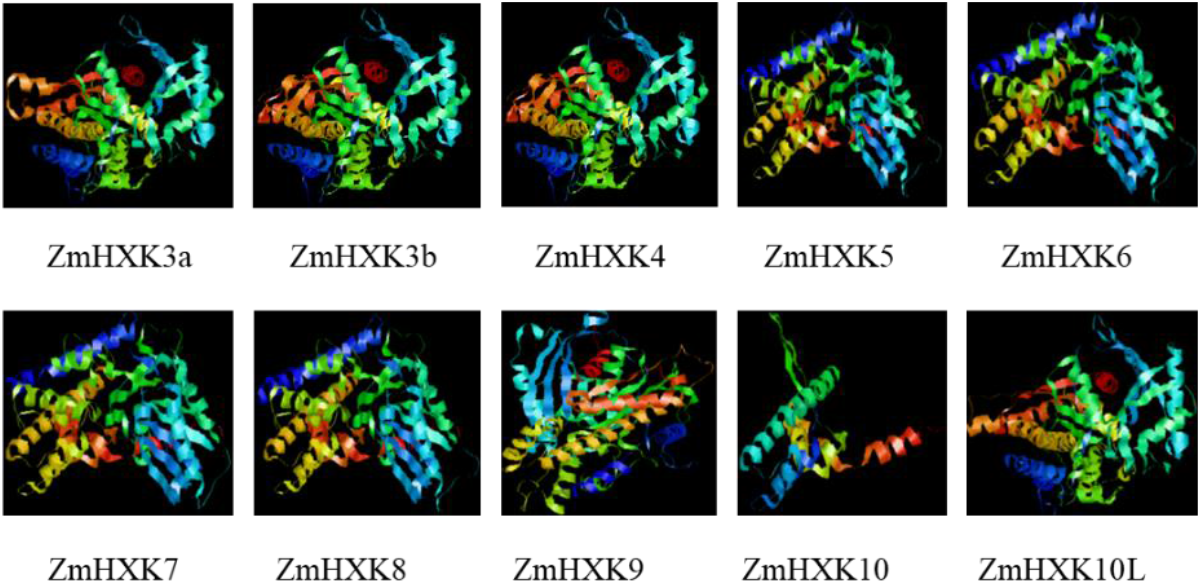
The three-dimensional modeling of the all the Hexokinase proteins in Oryza sativa and maize.

**Fig. 6.**
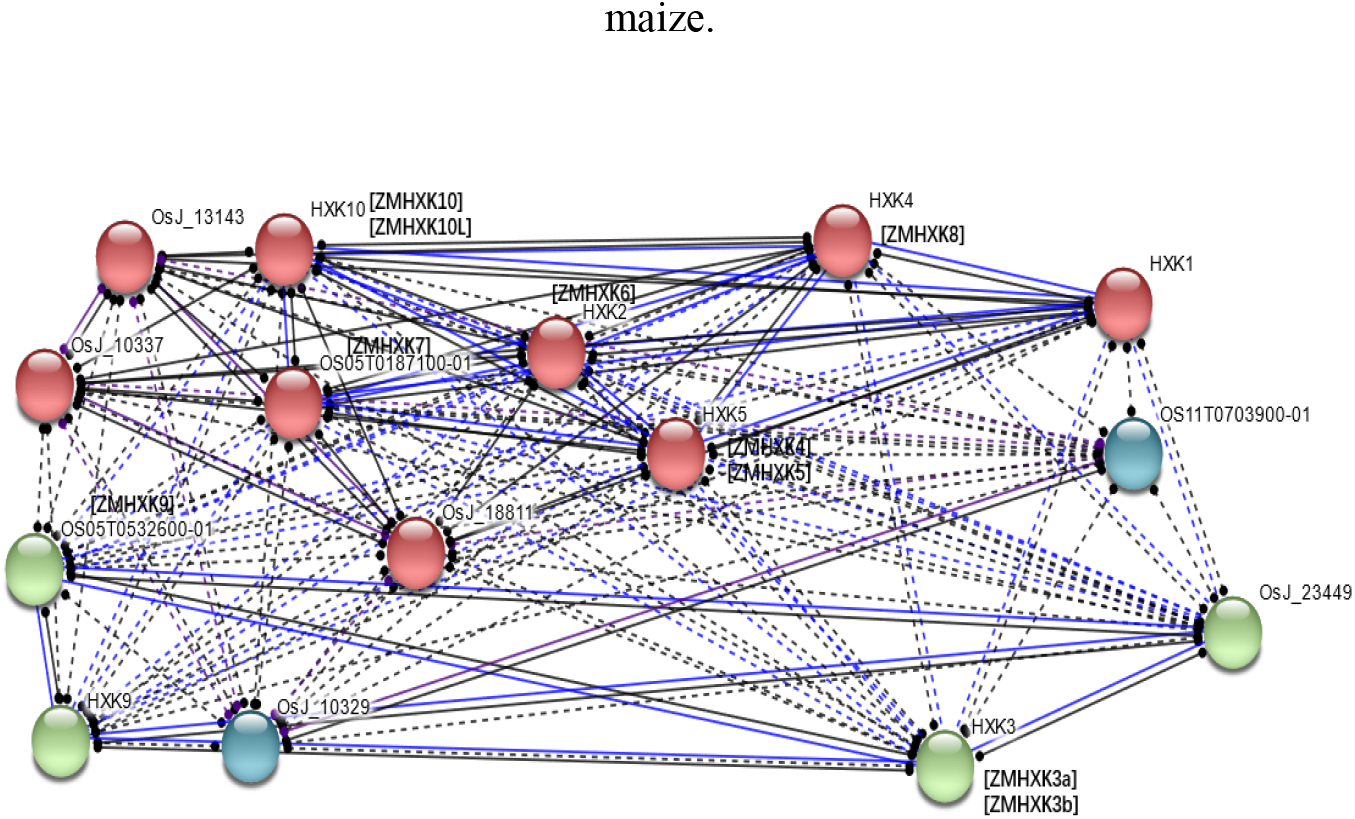
The prediction of the interation network of Hexokinase proteins in maize and Oryza sativa based on the interactions of their orthologs in Arabidopsis.

### 3.6 Expression Analysis of HXK Gene Family in Roots and Leaves under Drought or Salt Stress

To gain insight into potential functions, quantitative RT-PCR was employed to determine the expression patterns of hexokinase genes in leaves and roots of maize B73 line (Fig.7).

**Fig. 7.**
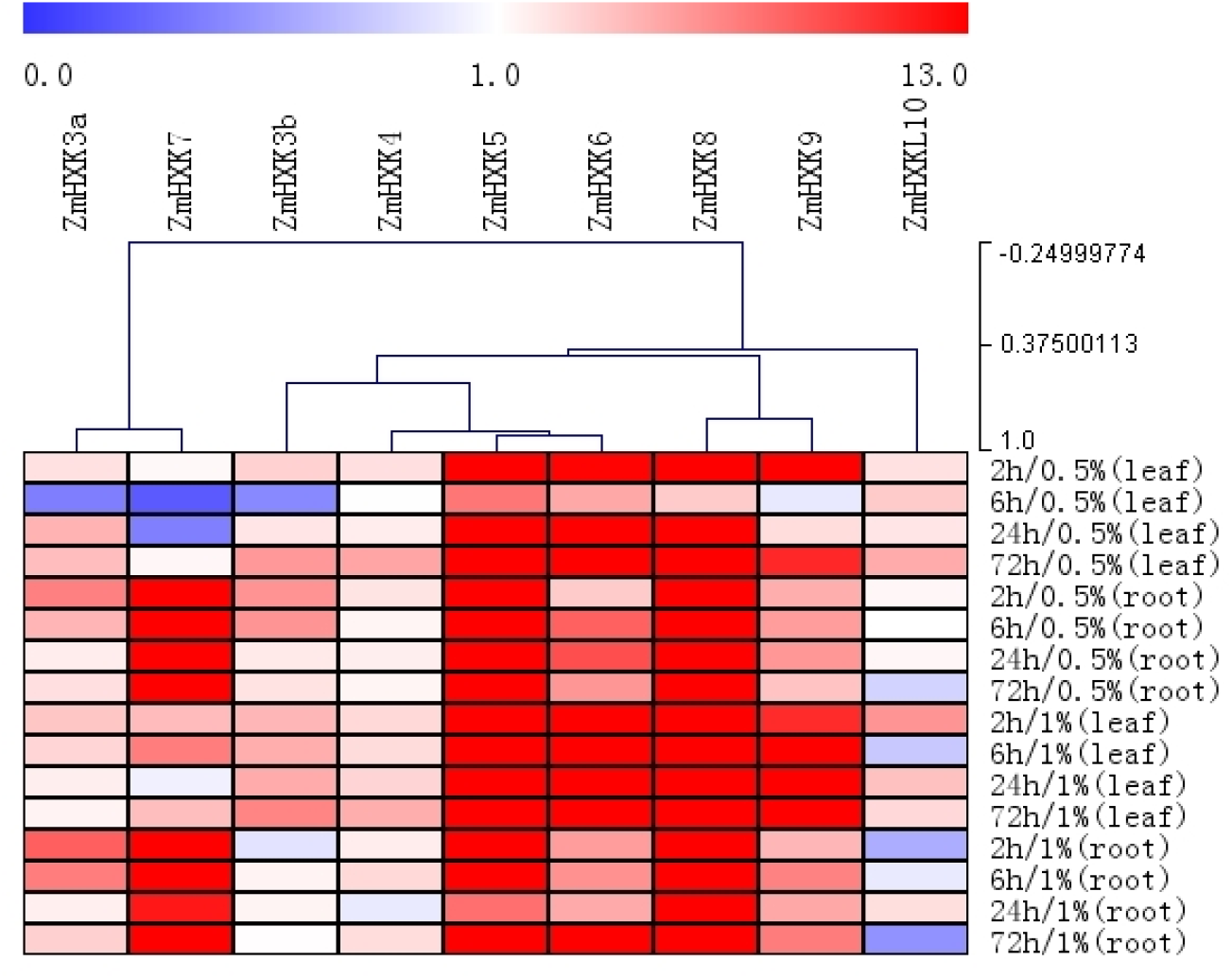
The relative expression of ZmHXK under salt stress treatment in Maize B73

Under both 0.5% and 1% concentration of NaCl treatment, the relative expression level of all genes in the leaves was upregulated at 2h in roots and leaves except for *ZmHXK10L* and *ZmHXK3b* in roots, which was down-regulated. *ZmHXK5, ZmHXK6* and *ZmHXK8* was predominantly expressed in roots and leaves during 2 to 72 hours and heat peak at 72h after salt stress.

The relative expression level of all genes in leaves and roots was upregulated in maize B73 line under drought treatment. *ZmHXK5* and *ZmHXK8* had significant expression in leaves and roots compared with other genes of the family (Fig.8).

**Fig. 8.**
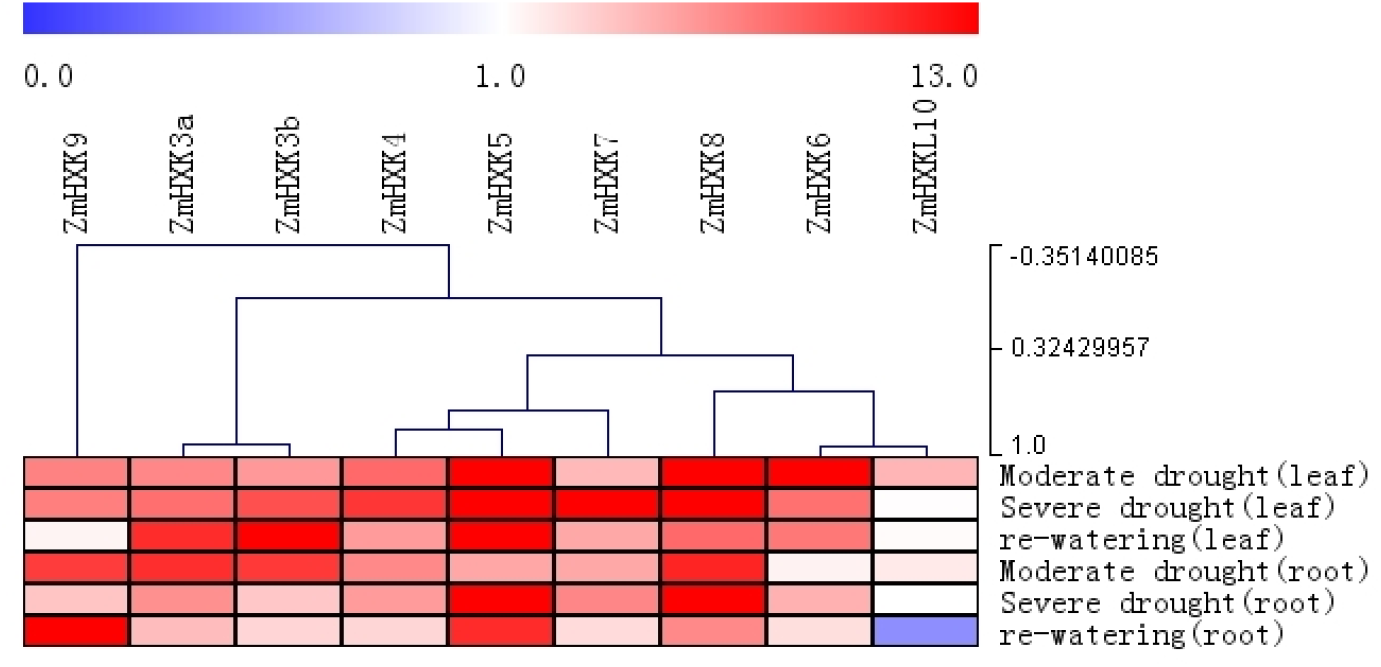
Relative expression of ZmHXK in maize B73 inbred line under drought stress

Similar to B73, the relative expression of most genes in the leaves and roots of Mo17 inbred lines was upregulated at 2h after salt treatment with 0.5% and 1% concentration of NaCl, except for *ZmHXK10L* at 1% salt concentration. The expression of *ZmHXK8* had the highest expression level in leaves with 1% NaCl at 72h(Fig.9).

**Fig. 9.**
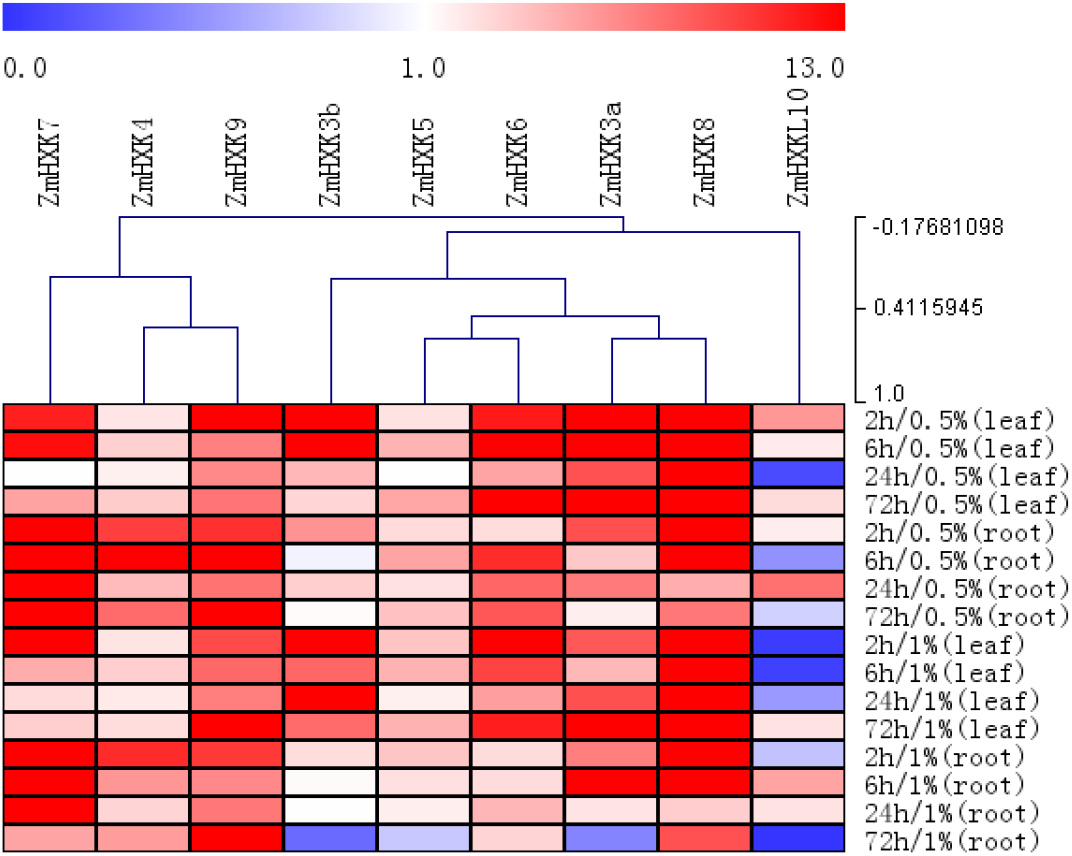
The relative expression of ZmHXK under drought stress treatment of maize Mo17

The mRNA of most genes in the leaves was higher compared with that in the roots, while that of *ZmHXK3a, ZmHXK5, ZmHXK7* and *ZmHXK8* in the root had remarkable expression under severe drought stress (Fig.10). The expression *ZmHXK6*,7,8,9,10 was downregulated after rewatering.

**Fig. 10.**
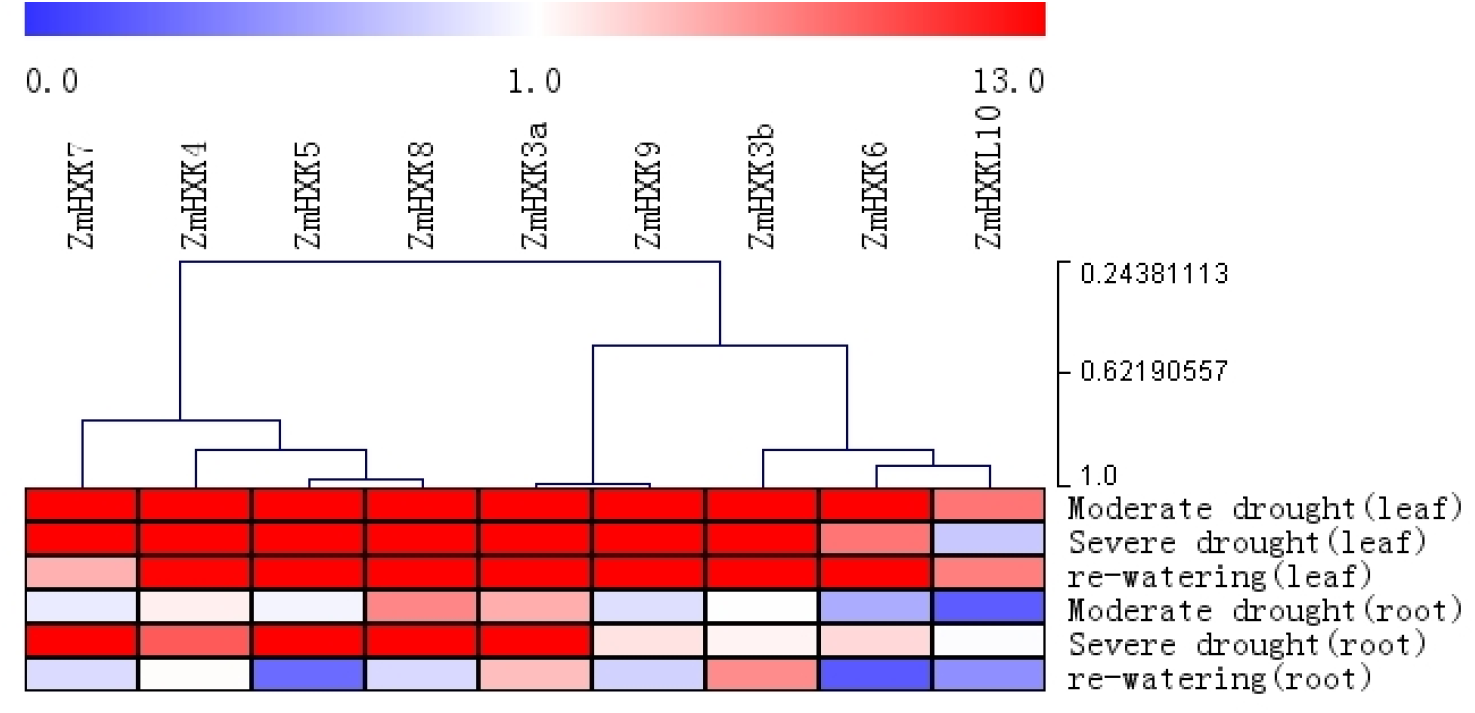
The relative expression of ZmHXK under drought stress treatment of rice Mo17

### 3.7 Molecular Characteristics of *ZmHXK7*

A full-length maize CDS fragment (GenBank Accession No: Zm00001d037689), without termination codon, was isolated by PCR with primers (FP:5’-CACCATGGAGAAGCAAGGCGTCGAC-3’;RP:5’-GTACTGCGAGTGGGCAGCT GCAATG −3’), using cDNA as a template, the PCR product was detected, and a 841bp fragment was obtained (Fig.11a). Then the PCR product was inserted into pGM-T vector and sequenced. Sequence analysis (BioXM 2.6) suggests that the fragment is identical with the known *ZmHXK7* gene. To investigate the function of *ZmHXK7* gene, four homozygous mutant *athxk3* strains were identified (Fig.11b) and labeled for use as subsequent transgenic receptors.

**Fig. 11.**
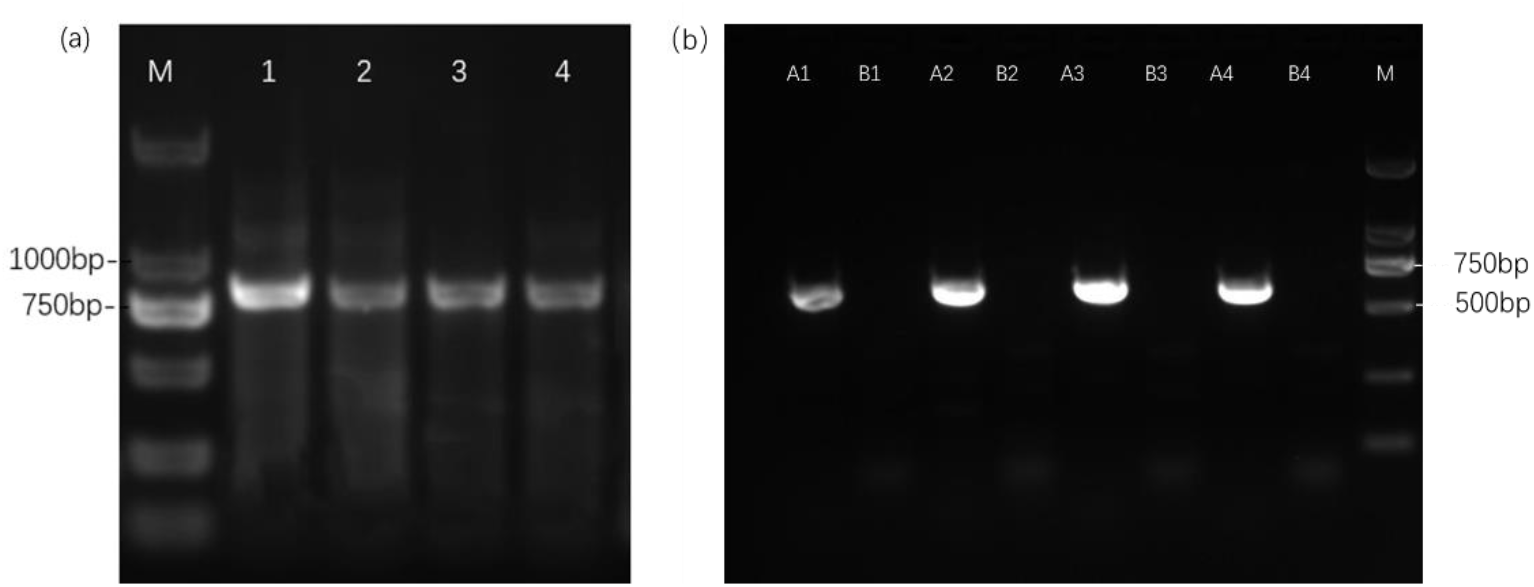
Target gene PCR amplification(a) and identification of homozygote of mutant *athxk3*(b) ((a): M: DL2000 Maker;1-4: ZmHXK7 target gene. (b): M: DL2000 Maker; A: BP+RP; B : LP+RP)

T3 seedlings for *athxk3* 35S:*ZmHXK7* were used for PCR analysis. The bands of about 1324bp were displayed (Fig12). The results showed that *ZmHXK7* genes of 16 seedlings except 3 and 4 were integrated into the genomes of WT and mutant Arabidopsis thaliana seedlings.

**Fig. 12.**
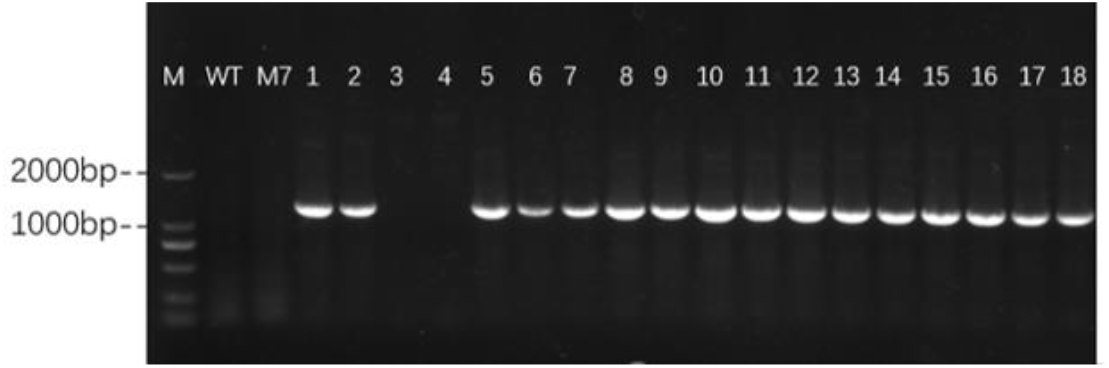
Molecular identification results of PCR (M: DL2000 Maker;1-18: Transgenic lines)

### 3.8 Subcellular localization of ZmHXK7 protein

As shown in (Fig.13), signals of GFP-ZmHXK7 were almost displayed in cytoplasm. For transgenic plants transformed with negative control pMDC45-GFP, GFP was expressed in the cytoplasm.

**Fig. 13.**
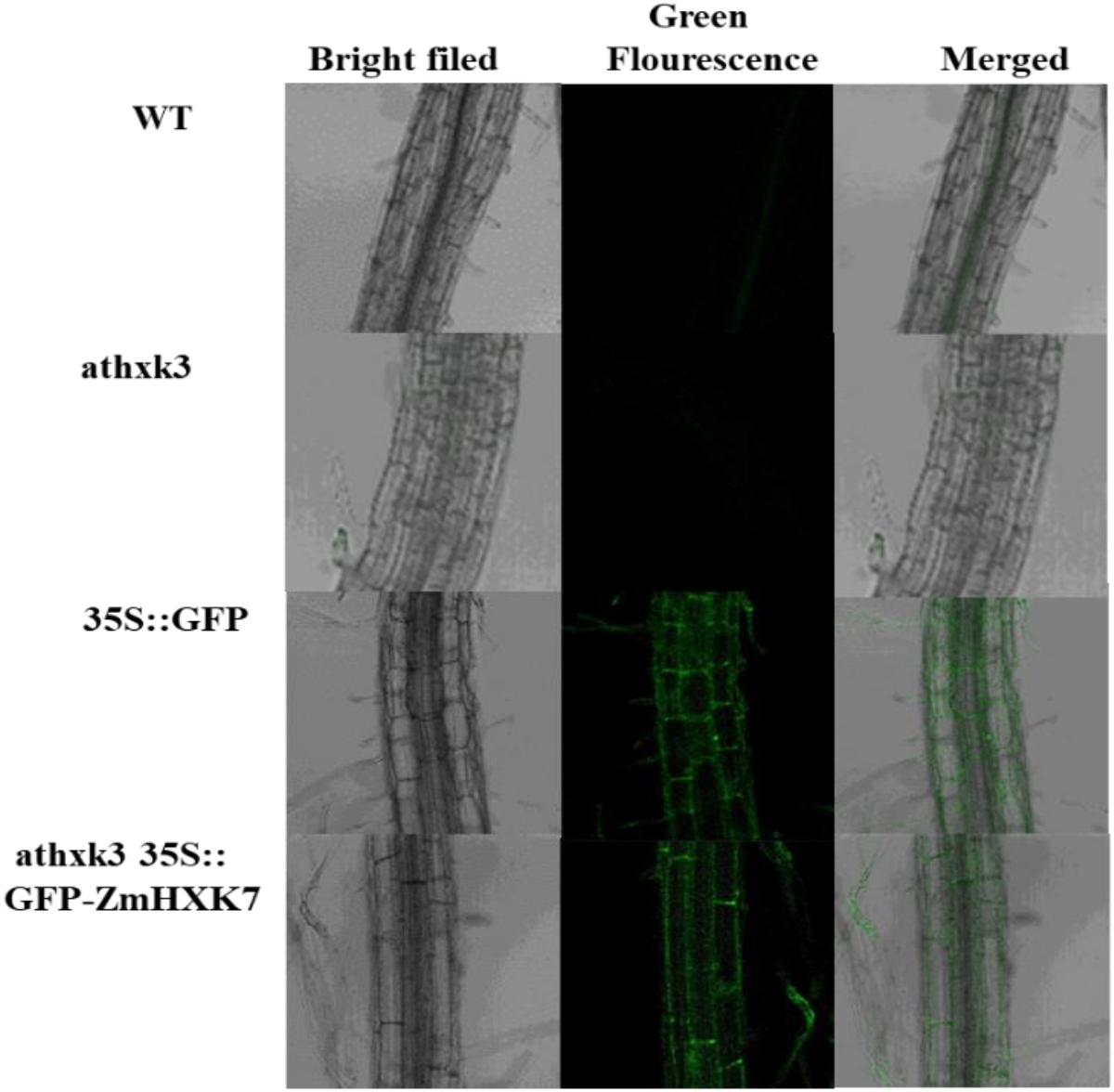
Subcellular localization of transgenic plant GFP-ZmHXK7

### 3.9 Western blot analysis of transgenic Arabidopsis plant

To examine ZmHXK7 protein expression, transgenic Arabidopsis seedlings *athxk3* 35S::*ZmHXK7* were further analyzed for ZmHXK7 protein integrated GFP via immunoblotting with antibody that was specific to GFP using Western blot. Our results show that transgenic Arabidopsis seedlings expressed the integrated protein with an expected molecular mass of 57KDa (Fig.14). Thus, the ZmHXK7 protein of the transgenic Arabidopsis in WT and mutant plants was successfully detected.

**Fig. 14.**
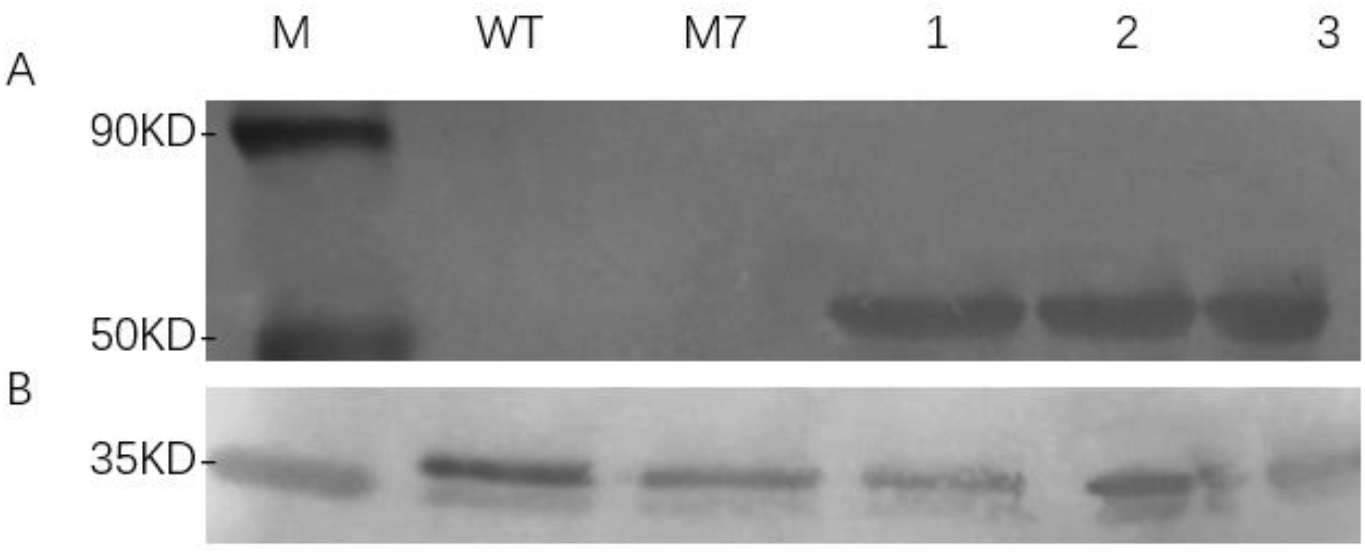
Protein expression analysis of transgenic lines (M: Protein Marker, WT: wild-type Arabidopsis thaliana, *athxk3* mutant Arabidopsis thaliana, 1-3 represents C45 transgenic line A: target Protein band,57KDa B: internal reference Protein band GAPDH.36KDa)

### 3.10 HK activity of transgenic Arabidopsis plant

The hexokinase activity of WT, *athxk3* and *athxk3* 35S::*ZmHXK7* was measured (Fig.15) The results showed that the hexokinase activity of mutant *athxk3* was significantly lower than that of WT, but the hexokinase activity of mutant *athxk3* was increased after transgenic *ZmHXK7* was complementary, even higher than that of WT, indicating that *ZmHXK7* was successfully expressed.

**Fig. 15.**
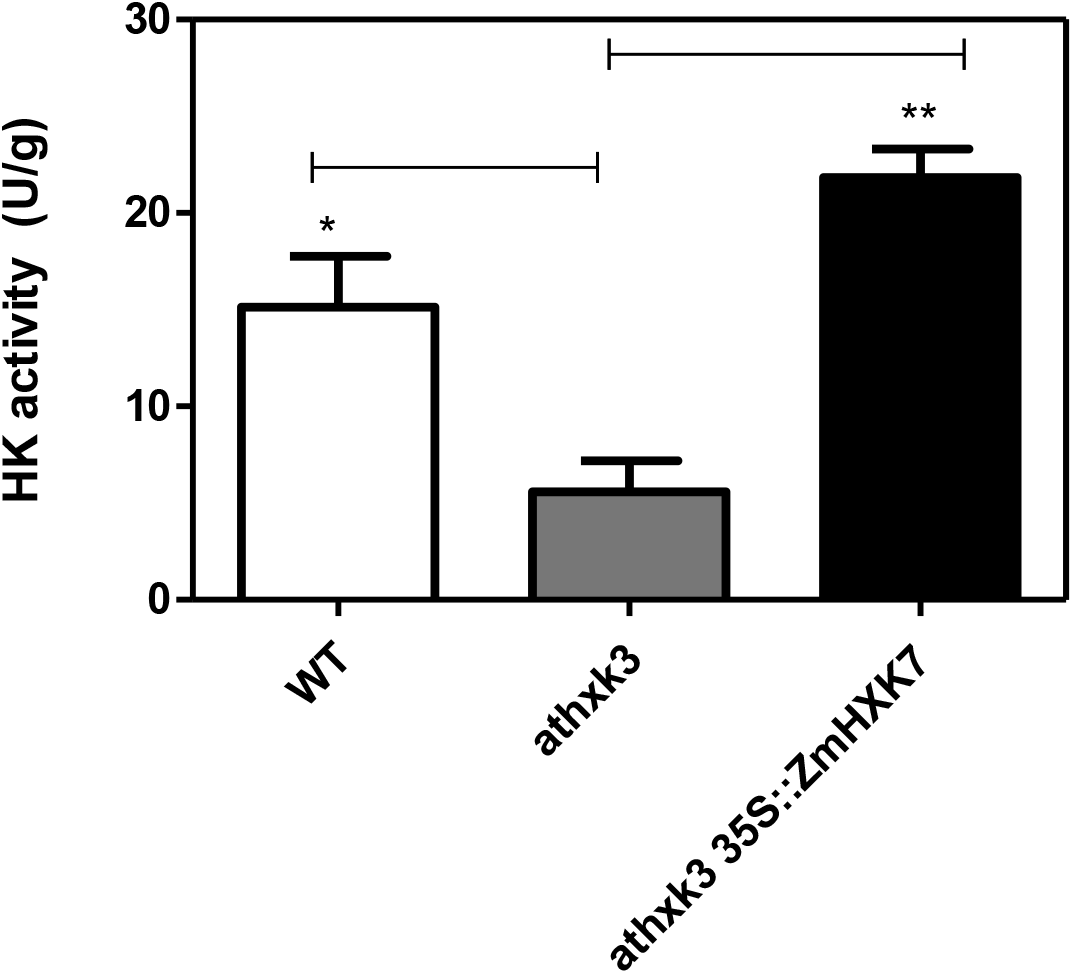
HK activity of WT, *athxk3, athxk3* 35S:: *ZmHXK7*. P<0.05 was considered significant *), P<0.01 was considered very significant(**)

### 3.11 Transgenic ZmHXK7 promotes plant stem growth

WT, *athxk3, athxk3* 35S::*ZmHXK7* grew in the MS medium for 15 days and then transplanted to vegetal soil for one month. We observed that the growth of leaf of WT, *athxk3* 35S::*ZmHXK7* lines was not grossly different, but growth of WT and *athxk3* 35S::*ZmHXK7* was much better than that of *athxk3*. Plant height of WT and *athxk3* 35S::*ZmHXK7* was higher than that of *athxk3* by 71.3% and 51.4% respectively(Fig. 16b). Complementary lines (*athxk3* 35S::*ZmHXK7*) had more branches, siliques and yield of seeds than *athxk3*(Fig.16c). There was no significant difference in the number of plant branches among the three groups. The growth of *athxk3* 35S::*ZmHXK7* lines recovered to a certain extent in comparison with *athxk3*.

**Fig. 16.**
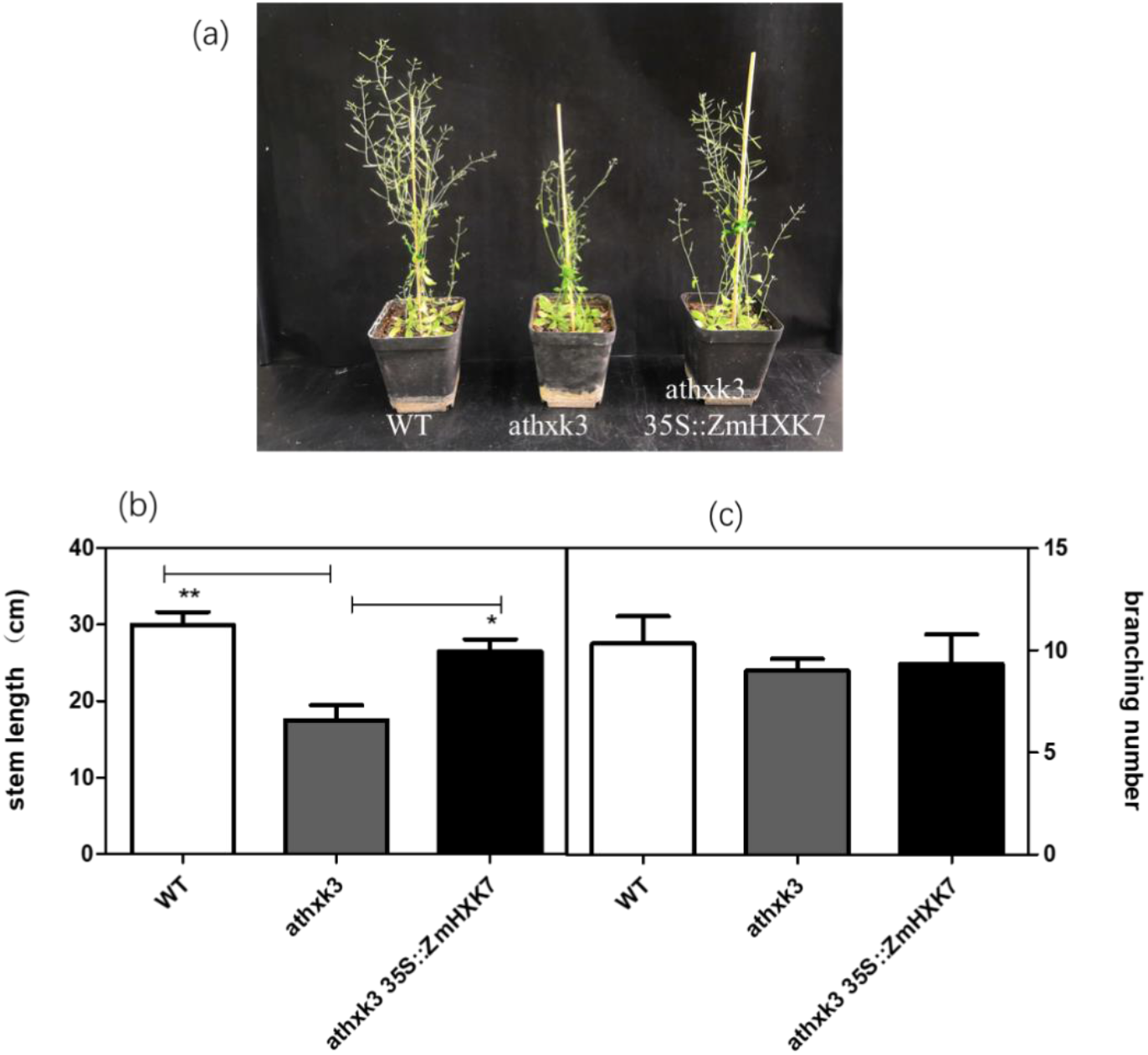
(a) The growth of WT, *athxk3* and T_3_ *athxk3* 35S::*ZmHXK7* seedlings. (b) The plant height of WT, *athxk3* and *athxk3* 35S::*ZmHXK7* of seedlings. (c) The number of branches of WT, *athxk3* and *athxk3* 35S::*ZmHXK7* seedlings. P<0.05 was considered significant (*), P<0.01 was considered very significant (**).

### 3.12 Exogenous glucose can partially inhibit the damage of salt to plant growth

WT, *athxk3* and *athxk3* 35S::*ZmHXK7* were grown on the MS plate supplemented with 0mM NaCl+0mM Glu, 150mM NaCl + 0mM Glu, 0mM NaCl+100mM Glu, 150mM NaCl+100mM Glu, respectively, and their growth is shown in the figure (Fig.17a).

**Fig. 17.**
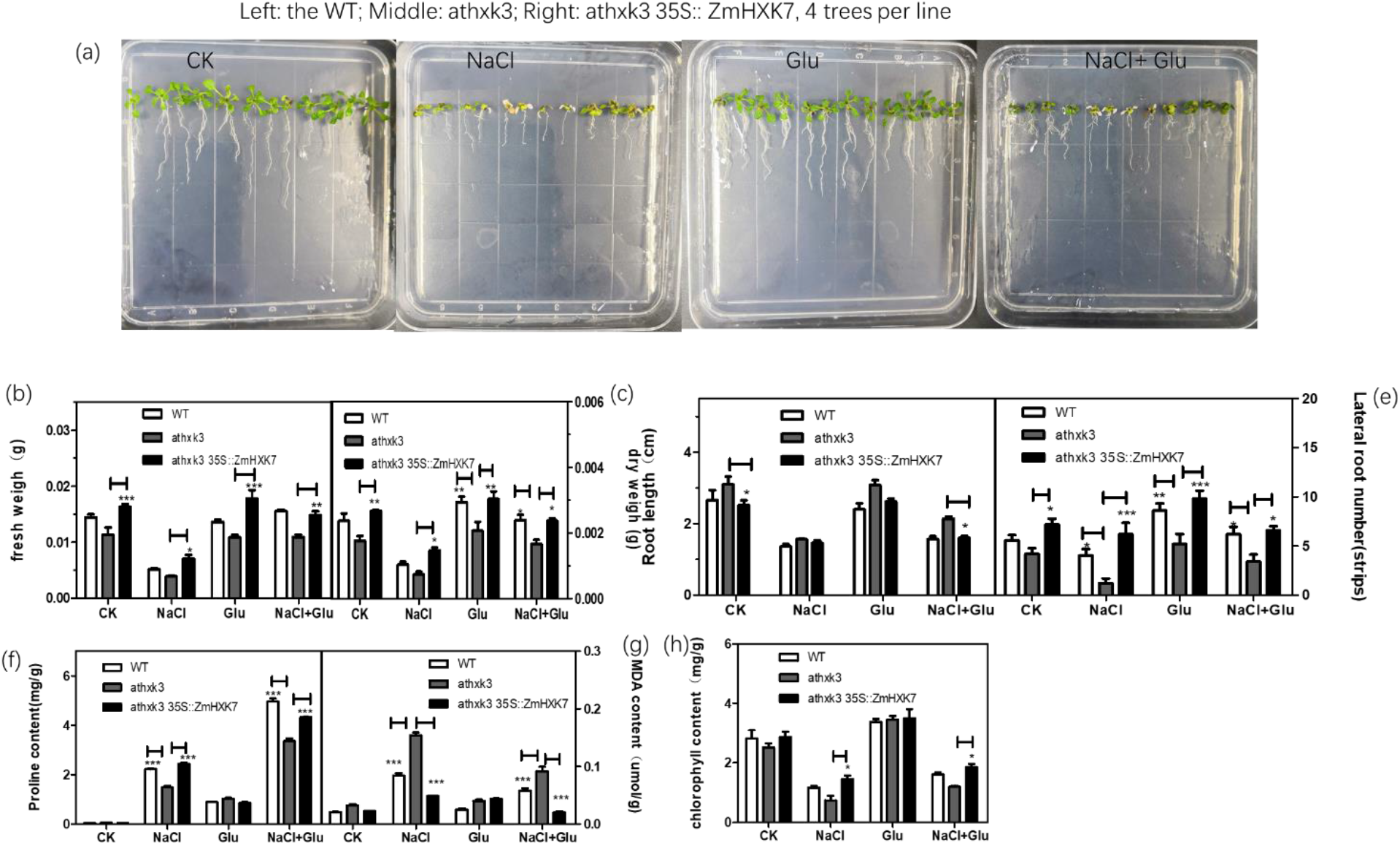
Exogenous glucose supplementation improves salt tolerance of Arabidopsis thaliana seedlings (a) Growth of WT, *athxk3, athxk3* 35S::*ZmHXK7* on salt and glucose plate (b) fresh weight of WT, *athxk3, athxk3* 35S:: *ZmHXK7* (c) dry weight of WT, *athxk3, athxk3* 35S::*ZmHXK7* (d) Proline content of WT, *athxk3, athxk3* 35S:: *ZmHXK7* (e) Chlorophyll content of WT, *athxk3, athxk3* 35S::*ZmHXK7* (f) root length of WT, *athxk3, athxk3* 35S:: *ZmHXK7* (g) Number of lateral roots of WT, *athxk3, athxk3* 35S:: *ZmHXK7* (h) Malondialdehyde content of WT, *athxk3, athxk3* 35S::*ZmHXK7* P<0.05 was considered significant(*), P<0.01 was considered very significant(**),and P<0.001 was considered extremely significant (***)

Salt treatment inhibited the growth of WT, *athxk3* and *athxk3* 35S::*ZmHXK7* under no glucose. Their fresh and dry weight of all above three groups decreased to some extent (as displayed in Fig.17b, c) under the salt stress. However, both the fresh and dry weight of *athxk3* 35S::*ZmHXK7* were increased in compared to *athxk3* under the same treatment. Exogenous glucose counteracts the effect on salt stress; the fresh weight and dry weight of WT, *athxk3* and *athxk3* 35S::*ZmHXK7* were increased compared to those at the 0 and 150mM NaCl stress.

Moreover, similar results were obtained with respect to chlorophyll and proline content (Fig.17d, e). These results indicate that the proline and chlorophyll contents of transgenic plants are significantly increased compared with atthxK3 under salt stress. Exogenous glucose improved plant salt tolerance. The proline and chlorophyll of WT, *athxk3* and *athxk3* 35S::*ZmHXK7* increased compared with that of 150mM NaCl stress, and the transgenic plant *athxk3* 35S::*ZmHXK7* was significantly higher than that of mutant *athxk3*.

Root length statistics under different treatments are shown (Fig.16f). The root length of *athxk3* was significantly higher than that of *athxk3* 35S::*ZmHXK7*, but the root length of *athxk3* was not significantly different from that of *athxk3* 35S:: *ZmHXK7* under salt treatment.

Lateral root number statistics under different treatments are shown (Fig.16g). The number of lateral roots of *athxk3* in different treatments decreased compared with WT, while the number of lateral roots of *athxk3* 35S::*ZmHXK7* was higher than that of *athxk3*. In control conditions, the number of lateral roots of *athxk3* 35S::*ZmHXK7* increased by 38.1% compared with that of *athxk3*. Under salt stress, *athxk3* 35S::*ZmHXK7* was 4.7 times that of athxk3.After glucose treatment, athxk3 35S::ZmHXK7 increased by 80.8% compared with *athxk3*; when treated with salt and glucose, C45 increased by 82.4% compared with *athxk3*. This indicates that the complementary line *athxk3* 35S::*ZmHXK7* can partially restore the growth of Arabidopsis thaliana compared with *athxk3*.

The content of malondialdehyde is an important index to detect the stress resistance of plants. A low content of malondialdehyde indicates a small degree of injury and a strong stress resistance under adverse conditions. The content of malondialdehyde (Fig.16h) was determined. Compared with the control group, the content of malondialdehyde decreased after salt treatment, and the content of malondialdehyde was lower than that of *athxk3*. However, the application of glucose under salt treatment inhibited the increase of MDA content under salt stress.

## 4 Discussion

As one of the most important crops in the world, *Zea mays* is sensitive to salt, and its growth and development are inhibited under salt stress (G 1977). It is urgent to cultivate salt-tolerant species of maize through biotechnology. Previous studies show that increased activity of HXK in guard cells will accelerate the closure of stomata, thereby reducing transpiration and reducing water loss in plants. In this current study, genome-wide ZmHXK gene family was characterized, and expression of 10 ZmHXK genes was analyzed in salt stress. mRNA of *ZmHXK7* increased after drought and salt treatment, and subsequent analysis found that overexpression of *ZmHXK7* improves the growth of Arabidopsis mutant plants under salt stress. These results suggest that ZmHXK will be a candidate gene for salt-tolerant transgenic breeding of plants in future.

In plants, glucose has emerged as a key regulator of many vital processes, including seedling development; root, stem, and shoot growth; photosynthesis; carbon and nitrogen metabolism; flowering; senescence; and stress responses (Kim et al. 2013). In the process, leaf starch of plant is synthesized during the day and degraded to glucose, maltose, or other component at night before it can be used(Niittyla et al. 2006). The glucose, once exported to the cytosol in the cytoplasm, can be phosphorylated by HXK to support sucrose synthesis, respiration and growth (Perata et al. 1997). In plant cells HXK not only promotes hexose phosphorylation, but also senses and conducts sugar signals (Olsson et al. 2003). In our study, the ZmHXK genes containing multiple cis-acting elements related to plant growth, implying that they not only play a regulatory role in photosynthesis, but also in the regulation of some cell metabolism of seedlings and the growth of mature plants. Moreover, *athxk3* 35S::*ZmHXK7* lines recovered phenotype of *athxk3* to a certain extent. Plant height of transgenic *athxk3* 35S::*ZmHXK7* increased compared with the *athxk3*, demonstrating that *ZmHXK7* is involved in development process. Corresponding with these results, *Athxk1* mutants and HXKI in tobacco induce delayed development and smaller leaves (Fox et al. 1998; Yu et al. 2013). *OsHXK7* overexpression promotes rice germination (Kim et al. 2016).

In the current study, different abiotic stress response elements, such as abscisic acid ABA response element (ABRE), methyl jasmonic acid response element, salicylic acid response element, ARE and GC-motif, DRE, MBS, LTR, TC-rich repeats, MYB, MYC and STRE, which are related closely to hormones and stress, appeared in *ZmHXK*. This shows that the ZmHXK gene plays an important role in response to abiotic stress. Overexpression of *ZmHXK7* increased the salt resistances of Arabidopsis mutant plants, further confirming that *ZmHXK7* is related in regulating the abiotic environment stress. Previous research showed that 4% exogenous glucose significantly improves the tolerance of apple seedlings to high salt stress (Sun et al. 2018), which is similar to our result, that application of exogenous glucose counteracted the effect on salt stress. Some papers indicate that HXK coordinates plant photosynthesis and transpiration by stimulating stomatal closure in guard cells (Galina et al. 1999; Shulga et al. 2009). For example, MdHXK1 gene was highly expressed induced by salt, low temperature and ABA(Kim et al. 2016),Similarly, the expressions of *ZmHXK5, ZmHXK7* and *ZmHXK8* were also significantly increased under salt and drought treatment. Furthermore, overexpressing *ZmHXK7* increased activity of HXK and improved the salt tolerance, suggesting that *ZmHXK7* modulates growth and development of plant in response to salt stress.

The primary root emerges from the seed during germination, and is very sensitive to environmental signals (such as gravity) (Gilroy and Masson 2007). As the root matures, the resting cells of its cycle begin to divide to form the lateral root primordia (Malamy 2008). and the lateral roots elongate and undergo further repeated branching (Kano-Nakata et al. 2013; Kano et al. 2011; Niones et al. 2012; YAMAUCHI et al. 1987a; YAMAUCHI et al. 1987b) to maintain root function, dry matter production and yield to a certain extent through water uptake and nutrient uptake. A positive role of HXK1 in lateral root development has previously been reported, as the number of lateral roots is reduced in gin2 (missing hxk1) mutants (Gupta et al. 2015). In the current study, like Athxk1, *athxk3* also decreased the number of lateral roots and its salt tolerance, while that of *athxk3* 35S::*ZmHXK7* was increased. Glucose also promotes root growth and lateral root formation at lower concentrations (Gupta et al. 2015; Malamy and Environment 2010; Ruan and Biology 2014; Ryan 2001). Here, exogenous glucose promoted lateral root growth of WT, *athxk3* 35S::*ZmHXK7*, with little effect on *athxk3*. The evidence shows that *ZmHXK7* may improve lateral root development by interacting with glucose. Auxin is reported to be related to lateral root formation, and mutants of auxin synthesis have lateral root defects(Negi et al. 2008). It is unknown that whether auxin in coordination with glucose signal regulate *ZmHXK7* expression to promote lateral root extending, and whether and how *ZmHXK7* or other HXK participate in stomatal closure. Detailed studies are required to understand of relationship between glucose sensing, activity of HXK and salt-tolerant phenotype such as stomatal closure to detect the molecular mechanisms that shed light on functions of ZmHXK in abiotic stress.

## Acknowledgements

This work was supported by the scientific and technological projects (172102110132) in Henan province, China.

## Author contributions

YQG and ZYT designed the research. QQL performed most of the experiments. All authors performed bioinformatic analyses, and collaboratively drafted the manuscript.

